# Context-dependent roles for autophagy in myeloid cells in tumor progression

**DOI:** 10.1101/2024.07.12.603292

**Authors:** Jayoung Choi, Gayoung Park, Steve Seung-Young Lee, Erin Dominici, Lev Becker, Kay F. Macleod, Stephen J. Kron, Seungmin Hwang

## Abstract

Autophagy is known to suppress tumor initiation by removing genotoxic stresses in normal cells. Conversely, autophagy is also known to support tumor progression by alleviating metabolic stresses in neoplastic cells. Centered on this pro-tumor role of autophagy, there have been many clinical trials to treat cancers through systemic blocking of autophagy. Such systemic inhibition affects both tumor cells and non-tumor cells, and the consequence of blocked autophagy in non-tumor cells in the context of tumor microenvironment is relatively understudied. Here, we examined the effect of autophagy-deficient myeloid cells on the progression of autophagy-competent tumors. We found that blocking autophagy only in myeloid cells modulated tumor progression markedly but such effects were context dependent. In a tumor implantation model, the growth of implanted tumor cells was substantially reduced in mice with autophagy-deficient myeloid cells; T cells infiltrated deeper into the tumors and were responsible for the reduced growth of the implanted tumor cells. In an oncogene-driven tumor induction model, however, tumors grew faster and metastasized more in mice with autophagy-deficient myeloid cells. These data demonstrate that the autophagy status of myeloid cells plays a critical role in tumor progression, promoting or suppressing tumor growth depending on the context of tumor-myeloid cell interactions. This study indicates that systemic use of autophagy inhibitors in cancer therapy may have differential effects on rates of tumor progression in patients due to effects on myeloid cells and that this warrants more targeted use of selective autophagy inhibitors in a cancer therapy in a clinical setting.

## Introduction

Macroautophagy (henceforth autophagy) plays crucial roles in cellular homeostasis^1^. Autophagy recycles essential metabolites, turns over damaged organelles, and clears intracellular aggregates that accumulate during aging, disease, and stress^2^. This catabolic pathway has been shown to play many important roles in tumor initiation and progression as well^3,4^. In one hand autophagy functions as *tumor suppressor* that prevents tumor initiation by removing genotoxic stress. On the other hand, autophagy can function as *tumor promoter* that helps established tumor cells to survive metabolic stress from their uncontrolled growth or therapeutic interventions. Consistently, chemical and genetic inhibition of autophagy was shown to inhibit the growth of established tumors and increase the killing effect of therapies^4,5^. Nevertheless, there are also contrasting results suggesting that autophagy may facilitate, rather than inhibit, the effectiveness of chemotherapy and radiation therapy^3,6^. More studies are necessary to elucidate the complex relationship between autophagy and various tumors in the context of tumor microenvironment, where tumor-stroma interactions as well as the nature of tumor itself play crucial roles in tumor progression^7,8^.

Tumors grow in close relationship with cells in stroma, such as fibroblast, endothelial cells, and immune cells^9^. Among the various cell types in the tumor microenvironment, macrophages have been known to play many crucial roles in controlling tumor progression^7,10^. Macrophages can be differentiated to multiple subtypes depending on their microenvironment^11^. Classically activated (also known as M1) macrophages express pro-inflammatory cytokines, chemokines, and effector molecules, which play crucial roles in immune defense against pathogens. In contrast, alternatively activated (also known as M2) macrophages rather produce anti-inflammatory and tissue remodeling substances, which are important for tissue repairing and homeostasis. In the tumor microenvironment, infiltrated monocytes differentiated to tumor-associated macrophages (TAM) with alternatively activated macrophage characteristics, and they promote tumor growth and metastasis in general^10^. Recent experimental and clinical evidence suggest a positive correlation of TAMs and the resistance of tumors to therapies and poor prognosis^12^. Thus, inhibiting the development and function of pro-tumor TAMs in the tumor microenvironment could be an effective anti-tumor strategy^10,12,13^.

Autophagy has been shown to play important roles in the differentiation and function of macrophages. The deletion of essential autophagy genes *Atg7* or *Atg16l1* in macrophages increases the secretion of pro-inflammatory cytokines (e.g. IL-1β, IL-6) and decreases the expression of anti-inflammatory cytokines (e.g. IL-10, TGF-β)^14,15^. Blocking of autophagic degradation by bafilomycin A_1_ or knock-down of an essential autophagy gene *Atg5* increase the expression of pro-inflammatory cytokines in M2 macrophages induced by hepatoma-cell-conditioned-media^16^. Autophagy is also activated and required for the M2 polarization of human and murine monocytes by colony stimulating factor 1 (CSF-1) and human peripheral blood monocytes by CCL2 and IL-6, prevalent cytokines in the tumor microenvironment^17,18^. Further, in a mouse model of obesity, autophagy-deficient macrophages are polarized to pro-inflammatory macrophages that induce immune responses^19^. Collectively, these *in vitro* and *in vivo* data suggest that the autophagy pathway may be required for the differentiation of macrophages to pro-tumor TAMs and consequently for the growth and metastasis of tumors.

To better understand the effect of autophagy inhibition on non-tumor cells in tumor progression, we investigated the effect of deleting critical autophagy genes in myeloid cells on autophagy-competent tumors using two mouse models, i.e., tumor implantation model and oncogene-driven tumor induction model. Our data demonstrate that autophagy inhibition in myeloid cells was sufficient to affect the progression of autophagy-competent tumors significantly; intriguingly, its effect on tumor progression was pro- or anti-tumor, depending on the context of tumor-stroma interaction. These data suggest that, as cell-intrinsic autophagy is known to suppress tumor initiation yet to support tumor progression, the autophagy status of non-tumor cells in the tumor microenvironment can also affect tumor progression positively or negatively, depending on the context of tumor-stroma interactions. Thus, systemic blocking of autophagy in both tumor and non-tumor cells, as following use of systemic autophagy inhibitors in cancer therapy, necessitates a context-dependent investigation of tumor progression for effective control of tumors at an organism level.

## Materials and Methods

### Mice

Mice from C57BL/6J background were used for implanted tumor studies. *Atg5^flox/flox^ +/- LysMcre, Atg7^flox/flox^ +/- LysMcre, and Atg16l1^flox/flox^ +/- LysMcre* mice were previously described^20^. *Atg14^flox/flox^ +/- LysMcre* were kindly provided by Dr. Shizuo Akira, Osaka University, Japan. MMTV-PyMT mice in FBV/NJ background were provided by Dr. Kay F. Macleod used for genetically induced tumor studies. To generate *MMTV-PyMT* +/- *Atg5^flox/flox^ +/- LysMcre* mice, *Atg5^flox/flox^ +/- LysMcre* mice were backcrossed into the FVB/NJ (The Jackson Laboratory) over 8 times, and then intercrossed with MMTV-PyMT mice. All mice were housed and bred at The University of Chicago under specific-pathogen-free conditions in accordance with federal and university guidelines.

### Tumor cell culture and implantation

MC38 colon carcinoma and B16.SIY melanoma (a derivative of B16-F10 expressing SIYRYYGL peptide)^21^ were provided by Dr. Yang-Xin Fu and Dr. Thomas F. Gajewski. The tumor cells were cultured in Dulbecco’s Modified Eagle Medium (DMEM; Corning) supplemented with 10 % heat-inactivated Fetal Bovine Serum (FBS; Biowest, US1520, Lot# 31A12), 100 U/ml penicillin and 100 μg/ml streptomycin (Corning). Cells were maintained at 20 ∼ 80 % confluency at 37°C and 5 % CO_2_ and used within 3 weeks once thawed from a frozen vial. Two days before the implantation, cells were seeded at 1x10^6^ cells per a 10 cm dish. On the day of implantation, cells were detached with Trypsin/Ethylenediaminetetraacetic Acid (EDTA) (Corning) and rinsed with complete media, followed by rinsing with 40 mL phosphate buffered saline (PBS) (Corning) at room temperature. Cells were re-suspended in 1 mL PBS and kept on ice until implantation. Live cells that were not stained with trypan blue were counted, and cell concentration was adjusted to 1x10^6^ cells / 1 mL PBS. 1x10^5^ viable cells in 100 uL PBS were implanted subcutaneously into the right flank of anesthetized mouse at 6∼8 week of age. To minimize variation due to cell death in PBS, the procedure from detaching cells to implantation was conducted within 1 hour. Tumor sizes were measured from day 5 to day 29 post implantation, and tumor volume (mm^3^) was calculated by multiplying the longest diameter, the shortest diameter, and the height of tumors.

The MMTV-PyMT induced tumor cells were isolated from 13-week-old MMTV-PyMT mice. Tumor tissues were dissociated into single cells by manual mincing using a dissecting scissors, followed by enzymatic digestion with dissociation solution: 2.5 mg/mL collagenase 4 (Worthington) and 200 μg/mL DNase I (Sigma-Aldrich) dissolved in DMEM containing 5% FBS. 50 mg tumor in 1 mL dissociation solution was incubated for 60 min at 37°C with gentle shaking. Enzyme activities were quenched by adding 40 μL of 0.5 M EDTA (Sigma-Aldrich) in 1mL dissociation solution. Digested tissues were minced with syringe plunger and filtered through 70 µm nylon strainers (Falcon), and cells were washed with DMEM containing 5 % FBS. The dissociated cells from tumor tissue were cultured in the media; DMEM, 5 % FBS, 100 U/mL penicillin, 100 μg/mL streptomycin, 2 mM L-glutamine (Corning), 1 mM Sodium Pyruvate (Corning), 1x MEM nonessential amino acid (Corning), 0.5 μg/mL hydrocortisone (Sigma-Aldrich), 5 μg/mL insulin (Sigma-Aldrich), 10 ng/mL Epidermal Growth Factor (Sigma-Aldrich), and 10 μg/mL gentamicin (Sigma-Aldrich). The tumor cells at passage 8 – 10 were used for subcutaneous implantation as described above.

### MMTV-PyMT breast tumor burden and lung metastasis

Tumor burden was determined by measuring mass (g) of breast tumors from each mouse at the age of 65, 80 and 95 days. Metastasis was measured by number of metastatic foci in lungs. Formalin fixed lungs were paraffin embedded, sectioned and stained with hematoxylin and eosin (H&E). The number of metastatic foci (>5 cells) were counted on 6 sections taken every 100 mm from each lung^22^.

### Flow cytometry analysis

Dissected tumor tissues were dissociated into single cell suspension by manual mincing using a dissecting scissors and enzymatic digestion with the dissociation solution for 60 min at 37°C in shaking incubator. Enzyme activities were quenched by adding EDTA to 20 mM. Digested tissues were minced with syringe plunger and filtered through 70 µm nylon strainers (Falcon) with DMEM containing 5 % FBS. Single cells were washed with flow media (PBS containing 1 % FBS and 1 mM EDTA). Cells were stained with Zombie NIR (BioLegend) to distinguish live and dead cells. Cells were washed with flow media and incubated with rat anti-mouse CD16/CD32 (TruStain FcX antibody, BioLegend) for 10 minutes on ice to prevent nonspecific antibody binding. Cells were washed in flow media and incubated for 30 minutes on ice with fluorophore-conjugated anti-mouse antibodies (BioLegend) to detect CD45 (30-F11), CD3 (17A2), CD4 (RM4-5), CD8 (53-6.7), CD25 (PC61), CD11b (M1/70), F4/80 (BM8), Ly-6G (1A8), or Ly-6C (HK1.4) using the manufacturers’ recommended or titrated concentrations. Intracellular staining with Foxp3 antibody (MF-14, BioLegend) was performed using Foxp3 / Transcription Factor buffer set (eBioscience). Data acquisition and analysis were performed using an LSR Fortessa system (BD Biosciences) and FlowJo version 9.2 software (Tree Star).

### Imaging of tumors

Tumor macrosection, immunofluorescence staining, imaging and analysis were conducted as previously described^23^. Briefly, tumor tissues were fixed with 2 % paraformaldehyde for 10 minutes at room temperature and embedded in 2 % agarose gel (LE Quick Dissolve Agarose, GeneMate) in 24 well plate. The embedded tumor tissues were sectioned by vibrating microtome (VT1200S, Leica). The macrosections were blocked with 10 mg/mL bovine serum albumin (BSA) and 10 % goat serum at 4°C for 1 hour, and stained with anti-mouse CD4 (GK1.5), CD8 (2.43) and F4/80 (CI:A3-1) antibodies (BioXCell). These antibodies were conjugated with fluorescent dyes: DyLight 594 NHS ester, DyLight 550 NHS ester and DyLight 680 NHS ester before the use. The stained macrosections were imaged with a Leica SP8 confocal microscope, and images were analyzed using Fiji (http://fiji.sc/Fiji). The distribution of immune cells in tumor sections was measured using distance map function to the tumor section outlines.

### T cell depletion

Rat anti-CD4 (GK1.5), rat anti-CD8 (YTS169.4), and rat IgG2b anti-KLH isotype control (LTF2) antibodies from BioXCell were diluted in PBS and injected intraperitoneally at 200 μg per mouse. Antibodies were injected every 5 days starting at a day prior to the tumor cell implantation^24,25^. Depletion was confirmed by flow analysis of peripheral blood mononuclear cells.

### RNA-seq

Myeloid cells were isolated from tumor tissues using EasySep mouse CD11b positive selection kit II (STEMCELL TECHNOLOGIES), and tumor associated macrophages (TAMs, CD11b^+^ F4/80^+^ Ly6G^-^ Ly6C^-^) were further isolated by Fluorescence Activated Cell Sorting (FACS). Total RNA from 0.5 – 1x10^6^ TAMs were extracted using RNA clean & concentrator kit (Zymo research). Novogene (www.novogene.com) conducted Eukaryotic RNA-seq and analysis. RNA-seq procedures were as follows: RNA quality test with Agilent 2100, library preparation using NEBNext ultra RNA library prep kit, and sequencing of 20M raw reads per sample on Illumina platform. Mapping reads to mouse reference genome was accomplished using STAR software. Gene expression level was determined by FPKM (the expected number of Fragments Per Kilobase of transcript sequence per Millions base pairs sequenced)^26^. Differential expression was analyzed using the DESeq2 R package^27^, and the P values were adjusted using the Benjamini and Hochberg’s approach for controlling the false discovery rate. The threshold of differential expression genes was padj < 0.05. Differentially expressed genes were analyzed for Gene Ontology using Metascape (http://metascape.org), and heatmaps were generated for the genes included in the top 5 gene ontology. The accession number for the RNA-seq data reported in this paper are GSE270910 (**Fig. 2G**) and GSE270912 (**Fig. 6A**).

**Figure 1.**
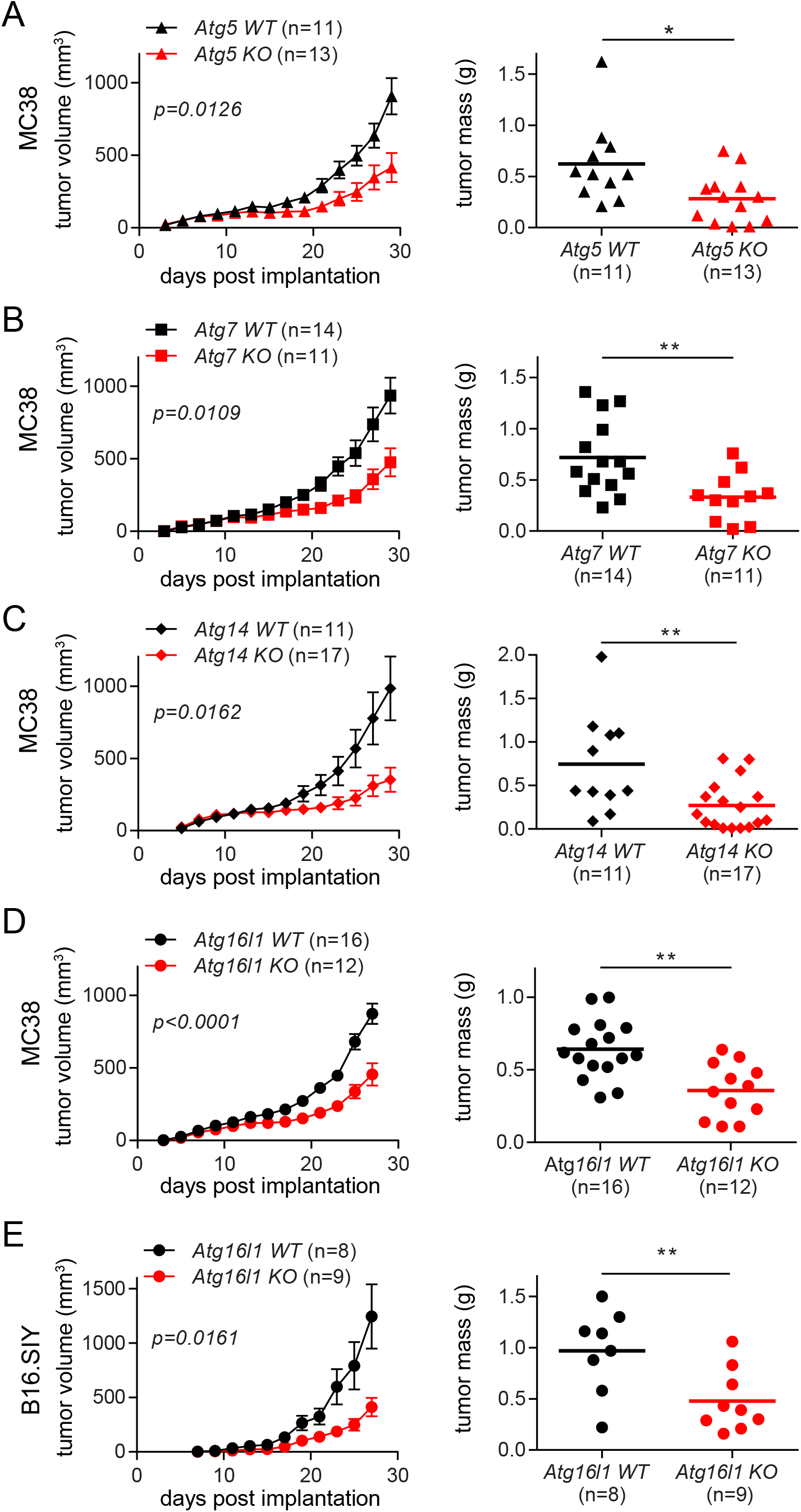
Autophagy deficiency in myeloid cells suppresses the growth of implanted tumors. (A-D) Tumor growth and tumor mass in mice with various autophagy genes deleted myeloid cells. 1x10^5^ MC38 cells were subcutaneously injected onto female mice. Tumor volumes on the left panel were measured every 2 days. At the end point, tumors were collected, and the weights were measured as shown on the right panel. Total number of mice used for each study (n) are indicated in each figure, and each data point in tumor mass represents each mouse. The *p* values for tumor volumes were calculated with two-way ANOVA, and the statistical significance for tumor mass were calculated with Mann-Whitney test. *p < 0.05, **p <0.01. (A) *Atg5 WT* is *Atg5*^flox/flox^ *and Atg5 KO* is *Atg5*^flox/flox^*+LysMcre.* (B) *Atg7 WT* is *Atg7*^flox/flox^ *and Atg7 KO* is *Atg7*^flox/flox^*+LysMcre.* (C) *Atg14 WT* is *Atg14*^flox/flox^ *and Atg14 KO* is *Atg14*^flox/flox^*+LysMcre.* (D) *Atg16l1 WT* is *Atg16l1*^flox/flox^ *and Atg16l1 KO* is *Atg16l1*^flox/flox^*+LysMcre.* (E) 1x10^5^ B16.SIY cells were subcutaneously injected onto female mice, and the same analysis as described above were performed.

**Figure 2.**
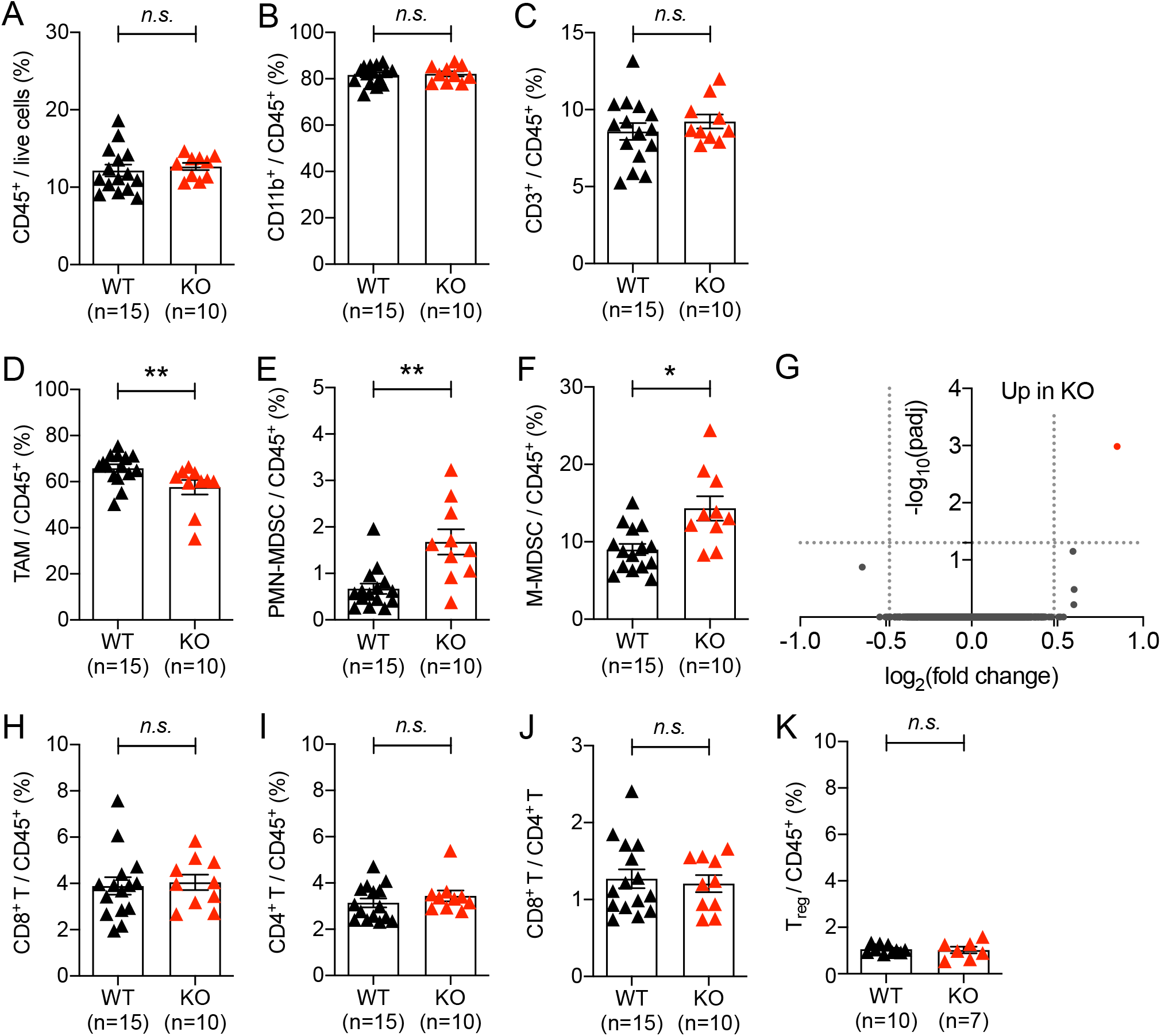
Impact of autophagy deficiency in myeloid cells on immune cell recruitment in implanted tumors. Tumor infiltrated immune cells were analyzed 21 days after 1x10^5^ MC38 cells were subcutaneously implanted into WT (*Atg5*^flox/flox^) and KO (*Atg5*^flox/flox^*+LysMcre*) mice. (A) Percentage of immune cells within the live cell population in tumor tissue. (B-F, H, I, K) Percentage of (B) myeloid cell (C) T lymphocyte (D) tumor associated macrophages, CD11b^+^ Ly6G^-^ Ly6C^-^, (E) polymorphonuclear MDSCs, CD11b^+^ Ly6G^high^ Ly6C^low^, (F) monocytic MDSCs, CD11b^+^ Ly6G^-^ Ly6C^high^, (H) CD8^+^ T cells, CD3^+^ CD8^+^, (I) CD4^+^ T cells, CD3^+^ CD4^+^ (K) regulatory T cells, CD3^+^ CD4^+^ CD25^+^ Foxp3+, within CD45+ population in tumor tissue. Total number of mice used for each study (n) are indicated in each figure, and each data point represents each mouse. Error bars indicate mean +/- SEM, and statistical significance was calculated with Mann-Whitney test. *p < 0.05, **p < 0.01, n.s.: not significant. (G) Gene expression from TAMs were analyzed by RNA-seq. Volcano plot shows the log_2_(fold change) in the expression of genes in TAMs from KO (*Atg5*^flox/flox^*+LysMcre*) mice compared to WT (*Atg5*^flox/flox^) mice. n=4 per group. Horizontal dashed line indicates the -log_10_ (padj) of 1.3 and vertical dashed line indicates the log_2_ (fold change) of +/- 0.48. (J) The ratio of CD8^+^ T cells to CD4^+^ T cells.

### Quantitative RT-PCR

Total RNAs prepared for RNA-seq were also analyzed for relative gene expression level in TAMs. cDNAs were synthesized using ImProm-II reverse transcriptase (Promega) with random hexamer. Quantitative PCR was conducted using SYBR green reagents in QuantStudio 3 Real-Time PCR system (Applied Biosystems). The primers used in this study are shown in Table S1.

### Statistical analysis

Data were plotted and analyzed with GraphPad Prism 7.0 software. Statistical significance was calculated with Mann-Whitney test, two-way ANOVA or unpaired T-test as indicated in the figure legends.

## Results

### Autophagy deficiency in myeloid cells suppresses the growth of implanted tumors

To investigate the effect of blocking autophagy in non-tumor cells on tumor progression, we examined the consequence of blocking autophagy in myeloid cells. Myeloid cells are known to engage at the site of tumor growth earlier than other immune cells and to play various pro-tumor functions^7,10^. We knocked out essential autophagy genes in myeloid cells through tissue-specific expression of Cre recombinase, *LysMcre*^28^. Autophagy genes are involved in multiple cellular pathways, other than canonical degradation through lysosomes^4,5^. Thus, we knocked out multiple autophagy genes essential for different pathways: induction of canonical autophagy (*Atg14*) and ubiquitin-like conjugation system (*Atg5*, *Atg7*, and *Atg16l1*)^29,30^. Using a syngeneic tumor implantation model, we examined whether autophagy deficiency in myeloid cells can affect the growth of autophagy-competent tumors. Upon subcutaneous injection of MC38 colon carcinoma cells into the flank of mice, we observed substantially slower growth of the implanted tumors in the mice with autophagy-deficient myeloid cells (i.e., *Atg5*^flox/flox^*+LysMcre, Atg7*^flox/flox^*+LysMcre, Atg14*^flox/flox^*+LysMcre,* and *Atg16l1*^flox/flox^*+LysMcre*), compared to their littermate control (i.e., *Atg5*^flox/flox^*, Atg7*^flox/flox^*, Atg14*^flox/flox^ and *Atg16l1*^flox/flox^) mice (**Fig. 1A-D**). Similar results were observed in both male and female mice, even though we analyzed and presented the growth of tumors separately based on sexes due to the significant difference in the rate of tumor growth between male and female mice (**Fig. S1**). The difference in tumor growths depending on the autophagy status of myeloid cells was not tumor-specific, because the same phenomenon was also observed upon implantation of B16 melanomas into the mice with autophagy-deficient myeloid cells and their littermate control mice (**Fig. 1E** **and S1. E**). Taken together, these data demonstrated that blocking autophagy in myeloid cells, the major immune cell type in tumor microenvironment, is sufficient to suppress the growth of autophagy-competent tumors.

### Autophagy deficiency in myeloid cells affects myeloid cell recruitment into implanted tumors

The implanted tumors in the mice with autophagy-deficient myeloid cells started growing differently compared to those in their littermate control mice at around 21 days-post-implantation (DPI), and the growth difference became significant by 28 DPI **(Fig. 1** **and S1)**. To understand this difference of tumor growth, we analyzed MC38 tumor-infiltrated immune cells at 21 DPI, when the growth difference began. The infiltration of total immune cells (CD45^+^), myeloid cells (CD11b^+^), and T cells (CD3^+^) were statistically indifferent between the two groups, WT (*Atg5*^flox/flox^) and KO (*Atg5*^flox/flox^*+LysMcre*) mice **(Fig. 2A-C**). However, there was a difference in the composition of infiltrated myeloid cells. The autophagy-deficient TAMs (CD45^+^ CD11b^+^ Ly6G^-^ Ly6C^-^) infiltrated tumors significantly less than autophagy-competent TAMs **(Fig. 2D)**; in contrast, the autophagy-deficient MDSCs (myeloid-derived suppressor cells), both PMN-MDSCs (polymorphonuclear MDSC, CD45^+^ CD11b^+^ Ly6G^high^ Ly6C^low^) and M-MDSCs (monocytic MDSC, CD45^+^ CD11b^+^ Ly6G^-^ Ly6C^high^), infiltrated into tumors significantly more than autophagy-competent MDSCs **(Fig. 2E-F)**. Although both TAM and MDSCs are known to promote tumor growth^31,32^, TAMs were the major myeloid cell population infiltrated into tumors in our model, accounting for over 50 % of CD45^+^ immune cell population **(Fig. 2D).** Thus, the decreased infiltration of TAMs may explain the slower growth of implanted tumors in the mice with autophagy-deficient myeloid cells.

The characteristics of TAMs, as well as their recruitment, could also be affected by autophagy deficiency. When we analyzed the total gene expressions in the recruited TAMs at 21 DPI via bulk RNA-seq, the entire transcriptomes were comparable between the autophagy-competent and -deficient TAMs (**Fig. 2G**). Expression of a single gene, *Lysozyme 1 (Lyz1)*, was significantly higher in the autophagy-deficient TAMs, possibly because of a compensatory upregulation due to the loss of *Lysozyme 2* in the LysM-cre mice used in this study^33^. Small differences in transcription between the two groups of TAMs might not be detected due to the choice of early time-point and relatively insensitive nature of bulk RNA-seq^34^. To complement this analysis, we performed quantitative PCR analysis using gene-specific primers for the representative markers of classically or alternatively activated macrophages and their key cytokines and chemokines (**Fig. S2**). Overall, consistent with the bulk RNA-seq data, most of those gene expressions did not differ significantly between the two groups, although a trend toward the pro-inflammatory side was noticed. In fact, the expression levels of a few pro-inflammatory genes (e.g., *Tnf, Saa3, S100a8*) were detected significantly more in the autophagy-deficient TAMs than the autophagy-competent TAMs (**Fig. S2B**). Collectively, these data suggest that the reduced number and the increased pro-inflammatory nature of the autophagy-deficient TAMs might contribute to the growth suppression of implanted tumor, in the mice with autophagy-deficient myeloid cells.

### Autophagy deficiency in myeloid cells suppresses the growth of implanted tumors by affecting T cell infiltration

To better understand the negative impact of autophagy-deficient myeloid cells on tumor progression, we examined the recruitment of T cells into the tumors at 21 DPI. The rate of tumor-infiltrated CD8^+^ T cells, total CD4^+^ T cells, and immune-suppressive CD4^+^ T_reg_ cells into whole tumors were comparable in those two groups of mice (**Fig. 2H-K**). In contrast, upon the quantitative spatial mapping of immune cells^23^, we observed a significant difference in the infiltration of immune cells within tumors (**Fig. 3**). In the mice with autophagy-competent myeloid cells, CD4^+^ and CD8^+^ cells were mainly located on the outer rim of tumors and F4/80^+^ macrophages infiltrated deeper into tumors (**Fig. 3A-D, S3**). In contrast, CD8^+^ cells infiltrated deeper into the center of tumors in the mice with autophagy-deficient myeloid cells, while CD4^+^ cells and F4/80^+^ macrophages localized similarly. These data suggest that a deeper penetration of CD8^+^ cells into tumors might contribute to the suppression of implanted tumor growth in the mice with autophagy-deficient myeloid cells.

**Figure 3.**
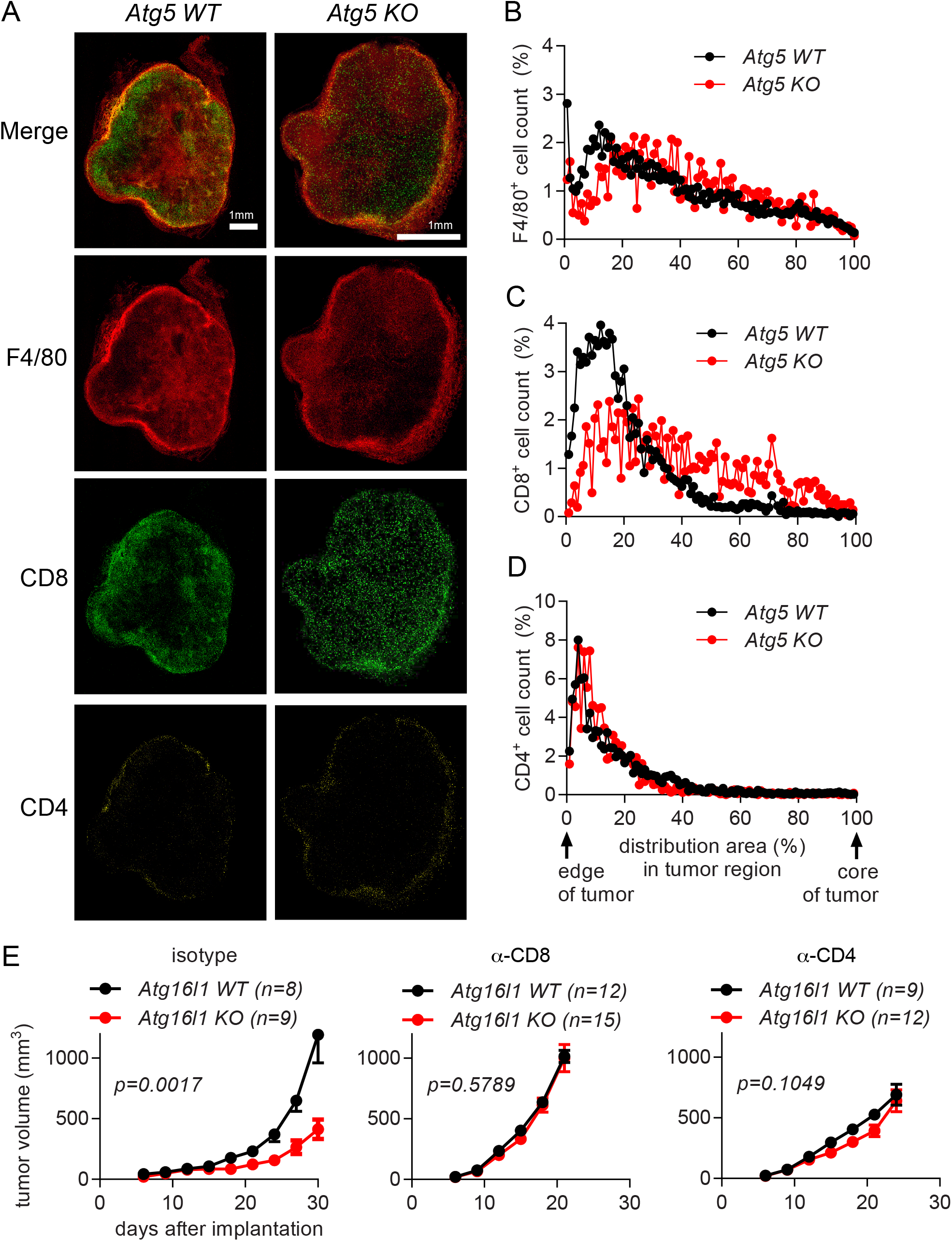
Autophagy deficiency in myeloid cells suppresses the growth of implanted tumors by affecting T cell infiltration. (A-D) Tumor tissues were collected at 21 days after 1x10^5^ MC38 cells were subcutaneously implanted into *Atg5 WT* (*Atg5*^flox/flox^) and *Atg5* KO (*Atg5*^flox/flox^*+LysMcre*) mice. n=3 per group. (A) Representative confocal microscopy images of the macrosections of tumors that were stained with anti-F4/80, anti-CD8 and anti-CD4 antibodies to visualize macrophages, CD8^+^ T cells and CD4^+^ T cells, respectively. The distribution of immune cells, (B) F4/80^+^ cells (C) CD8^+^ cells and (D) CD4^+^ cells, in tumor sections was measured from the edge of the tumor to the core of the tumor. (E) Tumor growth in *Atg16l1 WT* (*Atg16l1*^flox/flox^) and *Atg16l1* KO (*Atg16l1*^flox/flox^*+LysMcre*) female mice with T cell depletion. T cells were depleted by anti-CD8 or anti-CD4 antibodies. Isotype antibodies were treated as negative control. A day after antibody treatment, 1x10^5^ MC38 cells were subcutaneously implanted. Total number of mice used for each study (n) are indicated in each figure. Statistical significance was calculated with two-way ANOVA.

To examine whether the infiltrated CD4^+^ T and CD8^+^ T cells were responsible for the suppressed growth of tumors in the mice with autophagy-deficient myeloid cells, we depleted T cells and analyzed its effect on tumor growth. Antibodies to deplete T cells were injected before and throughout the tumor implantation, and the growth of tumors was measured (**Fig. 3E**). Depletion of either CD8^+^ T cells or CD4^+^ T cells removed the growth difference between tumors implanted to the mice with autophagy-competent or -deficient myeloid cells; such depletions enhanced the growth of tumors in the mice with autophagy-deficient myeloid cells to the level in the control mice, which was also enhanced especially upon CD8^+^ T cell depletion. Collectively, these data suggest that both CD4^+^ T and CD8^+^ T cells play important roles in the different growth of tumors in the mice with autophagy-competent or -deficient myeloid cells.

### Autophagy deficiency in myeloid cells promotes the growth of genetically induced tumors

Tumor implantation model is useful to investigate the role of myeloid cells in controlling the growth of established tumor cells^35,36^. To examine the role of autophagy-deficient myeloid cells in tumor generation and progression, we introduced mouse mammary tumor virus (MMTV)-polyoma middle T (PyMT) mouse model, which has been used to study mammary adenocarcinoma^37^. Mammary tumor develops upon mammary epithelium-specific expression of PyMT oncogene by MMTV long-terminal repeat promoter, followed by its metastasis to the lung. Early carcinoma can be detected in 9-week-old mice, and late carcinoma with lung metastasis can be measured in 13-week-old mice^38^. We generated *MMTV-PyMT+Atg5*^flox/flox^ (MP-WT) and *MMTV-PyMT+Atg5*^flox/flox^*+LysMcre* (MP-KO) mice and measured tumor mass at 65-day- (beginning of early carcinoma), 80-day- (early carcinoma) and 95-day- (beginning of late carcinoma) old mice. At 95 days, lungs were fixed and analyzed for metastasis. Tumor mass was indifferent between the MP-WT and MP-KO mice at 65 days; by 95 days, however, tumor growth was significantly promoted in the MP-KO mice **(Fig. 4A-C)**. Lung metastasis was also significantly increased in the MP-KO mice **(Fig. 4D).** These data indicate that autophagy-deficient myeloid cells promoted the growth and metastasis of genetically induced tumors, even though they suppressed the growth of implanted tumors **(Fig. 1** **and S1)**.

**Figure 4.**
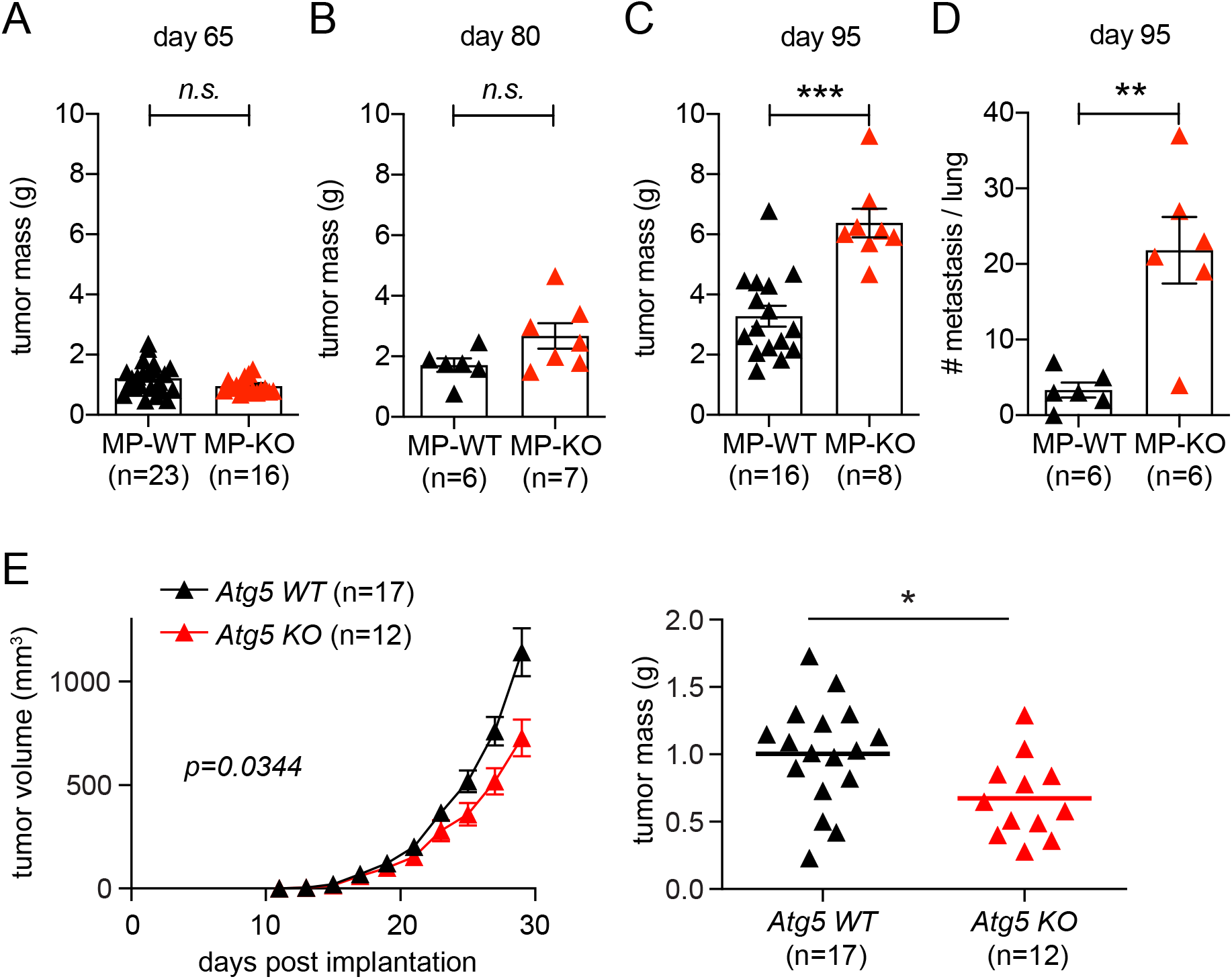
Autophagy deficiency in myeloid cells promotes the growth of genetically induced tumors while suppresses the growth of implanted tumors. (A-D) Mammary tumors were genetically induced in *MMTV-PyMT + Atg5*^flox/flox^ (MP-WT) and *MMTV-PyMT + Atg5*^flox/flox^ *+ LysMcre* (MP-KO) mice. Total mammary tumor mass per mouse was measured from (A) 65 day, (B) 80 day, and (C) 95 day old mice. (D) Each data point represents the average number of metastatic foci counted from 6 sections per lung. (E) 1x10^5^ tumor cells from MMTV-PyMT mice were subcutaneously implanted onto *Atg5*^flox/flox^ (*Atg5 WT*) and *Atg5*^flox/flox^ *+ LysMcre* (*Atg5 KO*) female mice. Tumor volumes on the left were measured every 2 days. At the end point, tumors were collected, and the weights were measured as shown on the right. Total number of mice used for each study (n) are indicated in each figure, and each data point in tumor mass represents each mouse. The *p* values for tumor volumes were calculated with two-way ANOVA and the statistical significance for tumor mass and lung metastasis were calculated with Mann- Whitney test. *p < 0.05, **p < 0.01, ***p <0.001, n.s.: not significant.

The differential effect of autophagy deficiency in myeloid cells on tumor growth in the autochthonous tumor model compared to the implanted tumor model could be due to the different context of interaction between myeloid cells and tumor cells. In the implantation model myeloid cells interact with already developed tumor cells; however, in this autochthonous model myeloid cells interact with tumor cells throughout their tumorigenesis and progression. To compare the impact of this context of interaction, we set up an implantation model using tumor cells from MMTV-PyMT mice. Late-stage carcinoma cells were cultured out of 13-week-old MMTV-PyMT mice and subcutaneously implanted onto the flank of syngeneic *Atg5*^flox/flox^ (*Atg5 WT*) and *Atg5*^flox/flox^*+LysMcre* (*Atg5 KO*) mice on a FBV/NJ background. Intriguingly, the implanted MMTV-PyMT tumor grew slower in the mice with autophagy-deficient myeloid cells (**Fig. 4E**), like the implanted MC38 and B16.SIY tumors (**Fig. 1** **and S1**). Taken together, these data demonstrate that autophagy-deficient myeloid cells can affect MMTV-PyMT tumor growth positively or negatively, depending on the context of interaction.

### Autophagy deficiency in myeloid cells affects CD8^+^ T cell recruitment and TAM activation in genetically induced tumors

To understand the tumor microenvironment at the stage of early carcinoma in the MMTV-PyMT model, we analyzed immune cells infiltrated into genetically induced tumor in 65-day-old mice. There was no significant difference in the infiltration of total immune cells (CD45^+^) and myeloid cells (CD11b^+^) in the tumors (**Fig. 5A, B**). In contrast to the tumor implantation model (**Fig. 2D-F**), we did not observe any significant difference in infiltrated TAM (CD45^+^ CD11b^+^ Ly6G^-^ Ly6C^-^), PMN-MDSCs (CD45^+^ CD11b^+^ Ly6G^high^ Ly6C^low^), and M-MDSCs (CD45^+^ CD11b^+^ Ly6G^-^ Ly6C^high^) between tumors from MP-WT and MP-KO mice **(Fig. 5C-E**). Further, the infiltration of CD4^+^ T cells and T_reg_ cells was also not affected by autophagy status in myeloid cells. However, we found significantly more CD8^+^ T cells infiltrated into the tumors in MP-KO mice compared to MP-WT mice (**Fig. 5F-I)**. It was previously shown that solid tumor burden at the stage of carcinoma formation in MMTV-PyMT mouse model is not affected by B and T cells in FVB/NJ background^22,39^. The pulmonary metastasis, however, is impacted by the pro-tumor properties of TAMs regulated by CD4^+^ T cells. Therefore, we further examined the characteristics of TAMs isolated from MMTV-PyMT tumors.

**Figure 5.**
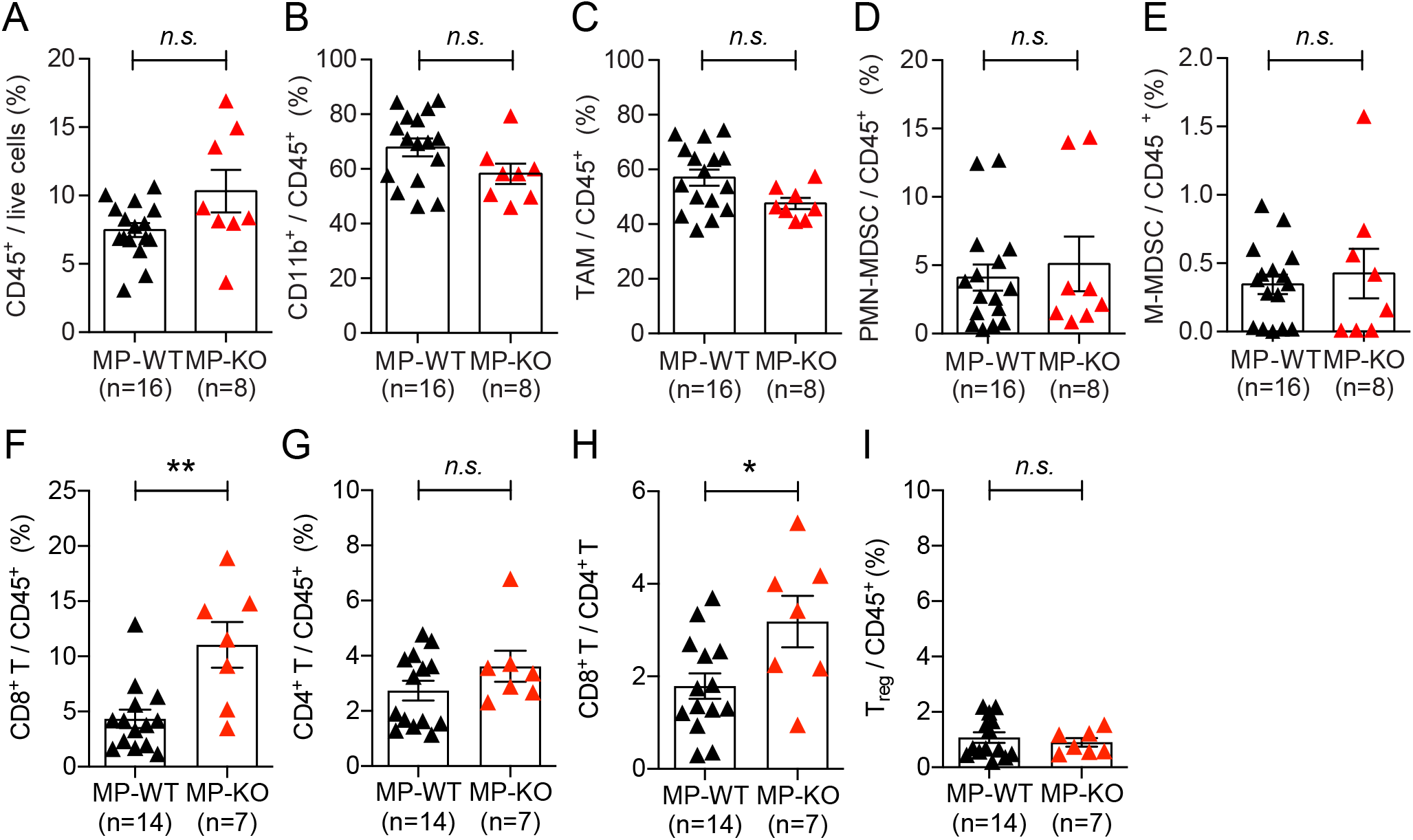
Impact of autophagy deficiency in myeloid cells on immune cell recruitment in genetically induced tumors. Tumors from 65-day-old mice, *MMTV-PyMT + Atg5*^flox/flox^ (MP-WT) and *MMTV-PyMT + Atg5*^flox/flox^ *+ LysMcre* (MP-KO), were collected, and infiltrated immune cells were analyzed. (A) Percentage of immune cells within the live cell population in tumor tissue. (B-E) Percentage of (B) myeloid cell (C) tumor associated macrophages, CD11b^+^ Ly6G^-^ Ly6C^-^, (D) polymorphonuclear MDSCs, CD11b^+^ Ly6G^high^ Ly6C^low^, (E) monocytic MDSCs, CD11b^+^ Ly6G^-^ Ly6C^high^ within CD45+ population in tumor tissue. (F,G,I) Percentage of (F) CD8^+^ T cells, CD3^+^ CD8^+^, (G) CD4^+^ T cells, CD3^+^ CD4^+^, (I) regulatory T cells, CD3^+^ CD4^+^ CD25^+^ Foxp3+ within CD45+ population in tumor tissue. (H) The ratio of CD8^+^ T cells to CD4^+^ T cells. Total number of mice used for each study (n) are indicated in each figure, and each data point represents each mouse. Error bars indicate mean +/- SEM, and statistical significance was calculated with Mann-Whitney test. *p < 0.05, **p < 0.01, n.s.: not significant.

When we analyzed the total gene expressions in the recruited TAMs in 65-day-old mice via bulk RNA-seq, the entire transcriptomes were comparable (**Fig. 6**); in both autophagy-competent and -deficient TAMs, genetic markers known for classically or alternatively activated macrophages were similarly expressed. Interestingly, autophagy-deficient TAMs in the tumors from MP-KO mice significantly up-regulated interferon response genes (**Fig. 6B, C**). In contrast, the genes related to blood vessel development and tissue repair were down-regulated (**Fig. 6B, D**). These data suggest that the different characteristics of the TAMs might have a significant effect on tumor progression and metastasis in the MMTV-PyMT induced tumor model.

**Figure 6.**
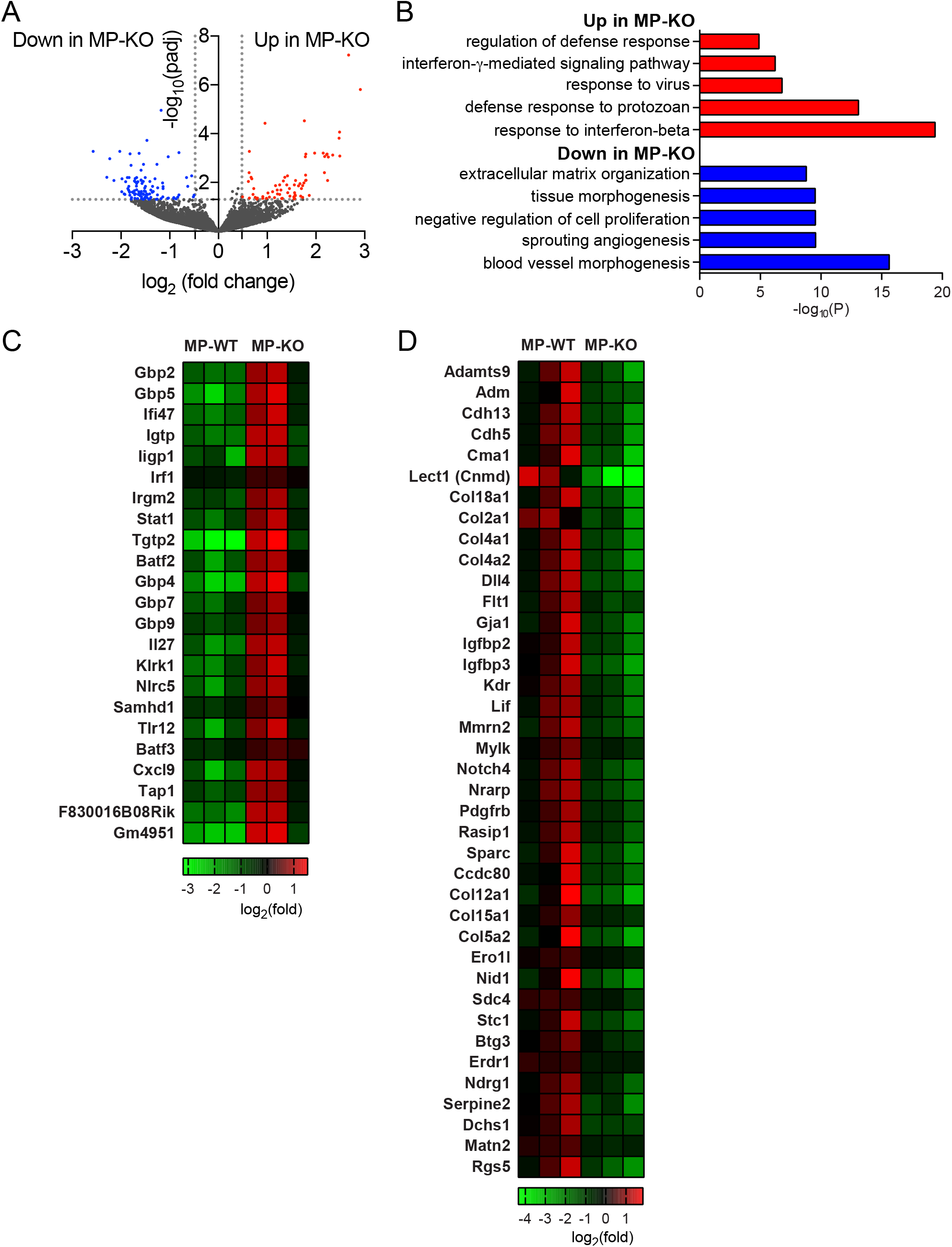
RNA-seq analysis of TAMs from genetically induced tumors. Tumors from 65- day-old mice, *MMTV-PyMT + Atg5*^flox/flox^ (MP-WT) and *MMTV-PyMT + Atg5*^flox/flox^ *+ LysMcre* (MP- KO), were collected and TAMs were isolated. Gene expression from TAMs were analyzed by RNA-seq. n=3 per group. (A) Volcano plot shows the log_2_(fold change) in the expression of genes in TAMs from the MP-KO mice compared to the MP-WT mice. Horizontal dashed line indicates the -log_10_ (padj) of 1.3 and vertical dashed line indicates the log_2_ (fold change) of +/-0.48. (B) Gene ontology of differentially expressed genes indicated as red and blue dots in the volcano plot (A). Top 5 gene ontology are depicted. (C) Heatmap showing expression level of genes that are up-regulated in MP-KO mice assigned to the top 5 gene ontology. (D) Heatmap showing expression level of genes that are down-regulated in MP-KO mice assigned to the top 5 gene ontology.

## Discussion

Cell-intrinsic autophagy plays dual roles in tumor progression, suppressing the transformation of normal cells into tumor cells while supporting the survival of established tumor cells^4^. Our study demonstrated that autophagy in non-tumor cells in tumor microenvironment also plays dual roles in tumor progression, depending on the context of cellular interactions. Autophagy in myeloid cells was required to *support* the growth of established tumors upon implantation; autophagy-deficient macrophages showed more proinflammatory characteristics and recruited more CD8+ T cells into the center of established tumors. In contrast, autophagy in myeloid cells was required to *suppress* the growth and metastasis of tumors upon genetic induction; autophagy-deficient macrophages in early carcinoma showed more interferon-activated and less tissue remodeling characteristics. Collectively, our data demonstrated that autophagy is important for tumor progression not only within tumor cells themselves but also in the surrounding immune cells.

Macrophages are versatile cells that play key functions to maintain homeostasis through the harmony of immune defense and tissue repair^7^. Within the tumor microenvironment, their phenotypic versatility and abundance impact many cell types and substantially affect tumor progression^40–42^. As a key process for cellular homeostasis, autophagy can affect the activation and differentiation status of macrophages substantially^12^. Induced autophagy promotes macrophage survival and accumulation in tumor environment^43^ and reduces the inflammasome activation and secretion of inflammatory cytokines^44–46^. Consistently, deletion of autophagy genes (e.g., *Atg5, Atg7, Atg14,* or *Atg16l1*) in myeloid cells leads to inflammasome activation and inflammatory IL-1β production^14^, polarization with increased pro-inflammatory and decreased anti-inflammatory gene expression^19^, and enhanced inflammation overall^47,48^.

Using the myeloid cell-specific perturbation system of autophagy (i.e., conditional deletion of key autophagy genes using *LysM-cre*), there have been multiple attempts to investigate the role of autophagy in non-tumor cells for tumor progression. For instance, Jinushi et al.^43^ subcutaneously injected B16 or MC38 tumor cells into *Atg5*^flox/flox^ and *Atg5*^flox/flox^*+LysMcre* mice and found no growth difference of tumors in those two groups of mice, until 17 days post-implantation. In contrast, upon intravenous injection of B16 or intraperitoneal injection of MC38 tumor cells, significantly less metastatic lesions were detected in *Atg5*^flox/flox^*+LysMcre* mice at 28 days or 21 days after injection, respectively. In the current study, subcutaneously implanted B16.SIY and MC38 tumors started growing differently in those two groups of mice at around 21 DPI, and the growth difference became significant by 28 DPI (**Figures 1** **and S1**). Thus, a difference in monitoring duration (i.e., 17 DPI vs 28 DPI) is likely to explain the different results. Nevertheless, the negative impact of autophagy-deficient myeloid cells on the progression of established tumor cells is consistent between the two studies.

Cunha et al.^49^ also made a similar observation that the growth of engrafted B16F10 melanoma, Lewis lung carcinoma, and MC38 adenocarcinoma are suppressed in *Atg5*^flox/flox^*+LysMcre* mice compared to *Atg5*^flox/flox^ mice. Intriguingly, they observed a similar tumor growth reduction in the mice with myeloid cells lacking genes required for both canonical autophagy and LC3-associated phagocytosis (LAP) (e.g., *Atg5, Atg7, Atg16l1*) but no difference in the ones lacking genes required only for canonical autophagy but not for LAP (e.g., *Atg14, Fip200, Ulk1*). In contrast to Cunha et al.^49^, we observed a significant growth reduction of MC38 tumors in *Atg14*^flox/flox^*+LysMcre* mice compared to *Atg14*^flox/flox^ mice, supporting a pro-tumor role of canonical autophagy, rather than LAP, in myeloid cells (**Fig. 1C, S1C**). This may be explained by a combination of differences in the engrafted tumor type (i.e., B16F10^49^ vs MC38), monitoring duration (i.e., 15 DPI^49^ vs 28 DPI), or the numbers of mice used (e.g., 7 mice^49^ vs 54 mice).

Interestingly, Cunha et al.^49^ further used a conditional mouse lung cancer induction model^50^ and found that the ablation of *Rubicon* in the hematopoietic compartment compromised tumor growth. Since *Rubicon* gene is required for LAP but not for canonical autophagy^51^, the authors conclude that LAP, but not canonical autophagy, in myeloid cells promotes the growth of genetically induced tumor. This is reminiscent of pro-tumor role of autophagy in cancer-associated fibroblast (CAF) in mammary tumor models driven by the PyMT oncogene^52^. The genetic loss of autophagy (*Atg5, Atg12*) in CAFs profoundly attenuates primary tumor growth and improve the survival of the tumor-bearing mice. In contrast, we observed the enhanced growth and metastasis of tumors in *MMTV-PyMT+Atg5*^flox/flox^*+LysMcre* (MP-KO) mice than *MMTV-PyMT+Atg5*^flox/flox^ (MP-WT) mice (**Figure 4**). Taken together, these studies demonstrate that the effect of autophagy and autophagy-related processes in neighboring cells on tumor progression should be examined in a context dependent manner.

The beneficial effects of systemic autophagy inhibition on tumor regression occur more rapidly than the detrimental metabolic and neurological responses, suggesting the possibility of an optimal therapeutic window for systemic autophagy inhibition as anticancer therapy^3–5^. However, the critical yet context-dependent role of autophagy in both tumor and non-tumor cells for tumor progression suggest that systemic therapeutic targeting of autophagy may have undesirable side effects^4^. In conclusion, this study supports careful investigation of effects of systemic autophagy blocking on tumor progression and development of more selective autophagy inhibitors for more effective control of tumors at the organism level.

## Acknowledgments

This work was supported in part by the Brinson Foundation Junior Investigator Grant, Cancer Research Foundation Young Investigator Award, an Institutional Research Grant (IRG- 58-004-53-IRG) from the American Cancer Society, the University of Chicago Comprehensive Cancer Center Support Grant (P30 CA14599), and Cancer Center Core facilities (DNA sequencing and genotyping, flow cytometry, integrated microscopy, and monoclonal antibody), the National Center for Advancing Translational Sciences of the National Institutes of Health (NIH UL1 TR000430), and NIH DP2 CA225208 to SH. SSL and SJK were supported by K99/R00 EB022636 to SSL and R01s CA199663 and CA232419 to SJK.

## Conflicts of Interest

All authors declare no conflict of interest.

## Author Contributions

Conceptualization: JC, SSL, LB, KFM, SJK, SH; Data curation: JC, GP, SSL, ED; Formal analysis: JC, GP, SSL; Funding acquisition: SH; Investigation: JC, GP, SSL, ED, SH; Methodology: JC, GP, SSL, KFM; Project administration: JC, SH; Resources: SJK, SH; Software: JC, GP, SSL, SH; Supervision: SJK, SH; Visualization: JC, SSL; Writing – original draft: JC, SH; Writing – review & editing: JC, GP, SSL, ED, LB, KFM, SJK, SH.

---

**Figure S1.**
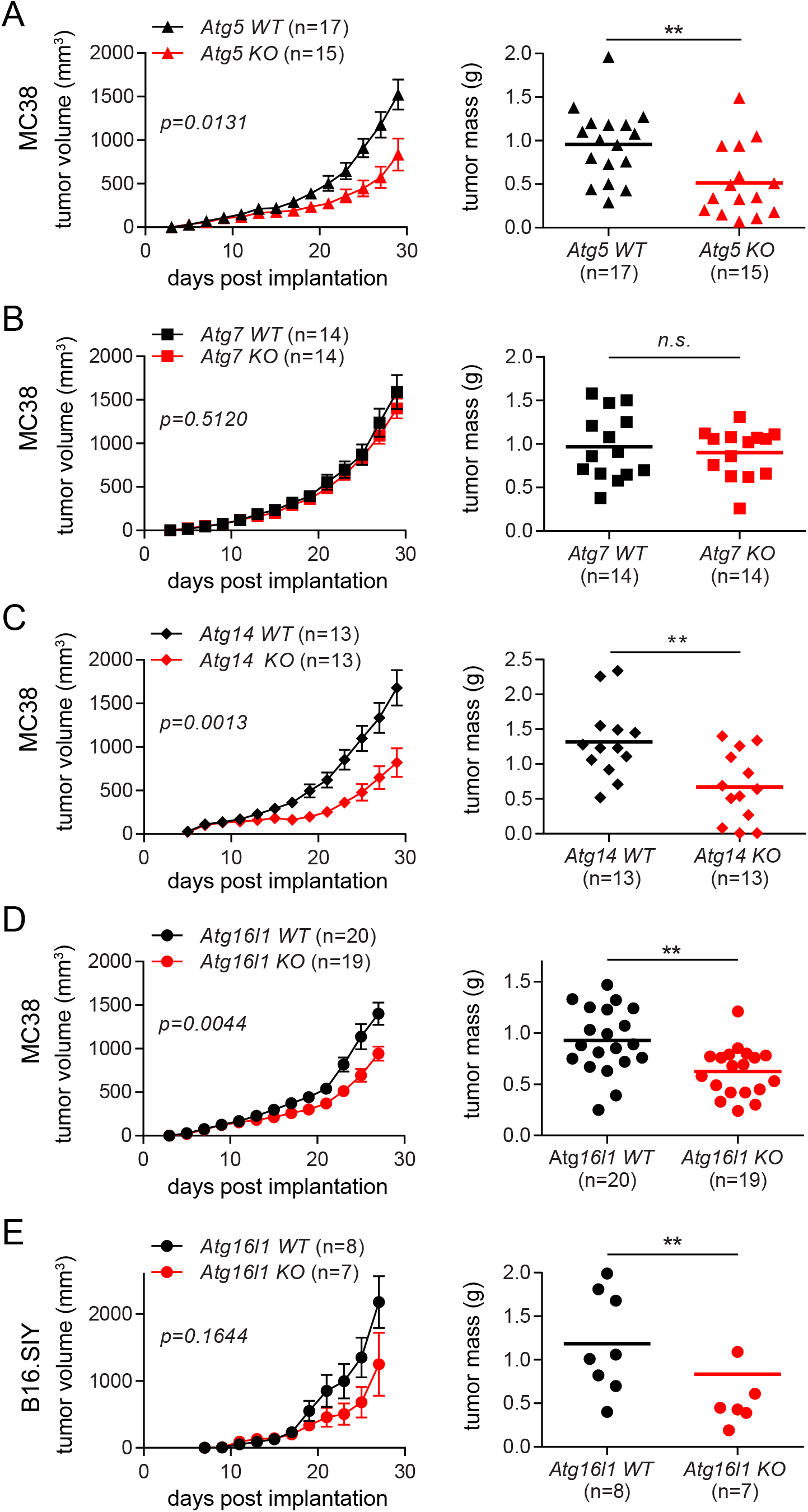
Autophagy deficiency in myeloid cells suppresses the growth of implanted tumors. (A-D) 1x10^5^ MC38 cells were subcutaneously injected onto male mice. Tumor volumes on the left panel were measured every 2 days for 27 – 29 days post implantation. At the end point, tumors were collected, and the weights were measured as shown on the right panel. Total number of mice used for each study (n) are indicated in each figure, and each data point in tumor mass represents each mouse. The *p* values for tumor volumes were calculated with two-way ANOVA and the statistical significance for tumor mass were calculated with Mann-Whitney test. *p < 0.05, **p < 0.005, n.s.: not significant. (A) *Atg5 WT* is *Atg5*^flox/flox^ *and Atg5 KO* is *Atg5*^flox/flox^*+LysMcre.* (B) *Atg7 WT* is *Atg7*^flox/flox^ *and Atg7 KO* is *Atg7*^flox/flox^*+LysMcre.* (C) *Atg14 WT* is *Atg14*^flox/flox^ *and Atg14 KO* is *Atg14*^flox/flox^*+LysMcre.* (D) *Atg16l1 WT* is *Atg16l1*^flox/flox^ *and Atg16l1 KO* is *Atg16l1*^flox/flox^*+LysMcre.* (E) 1x10^5^ B16.SIY cells were subcutaneously injected onto male mice, and the same analysis as described above were performed.

**Figure S2.**
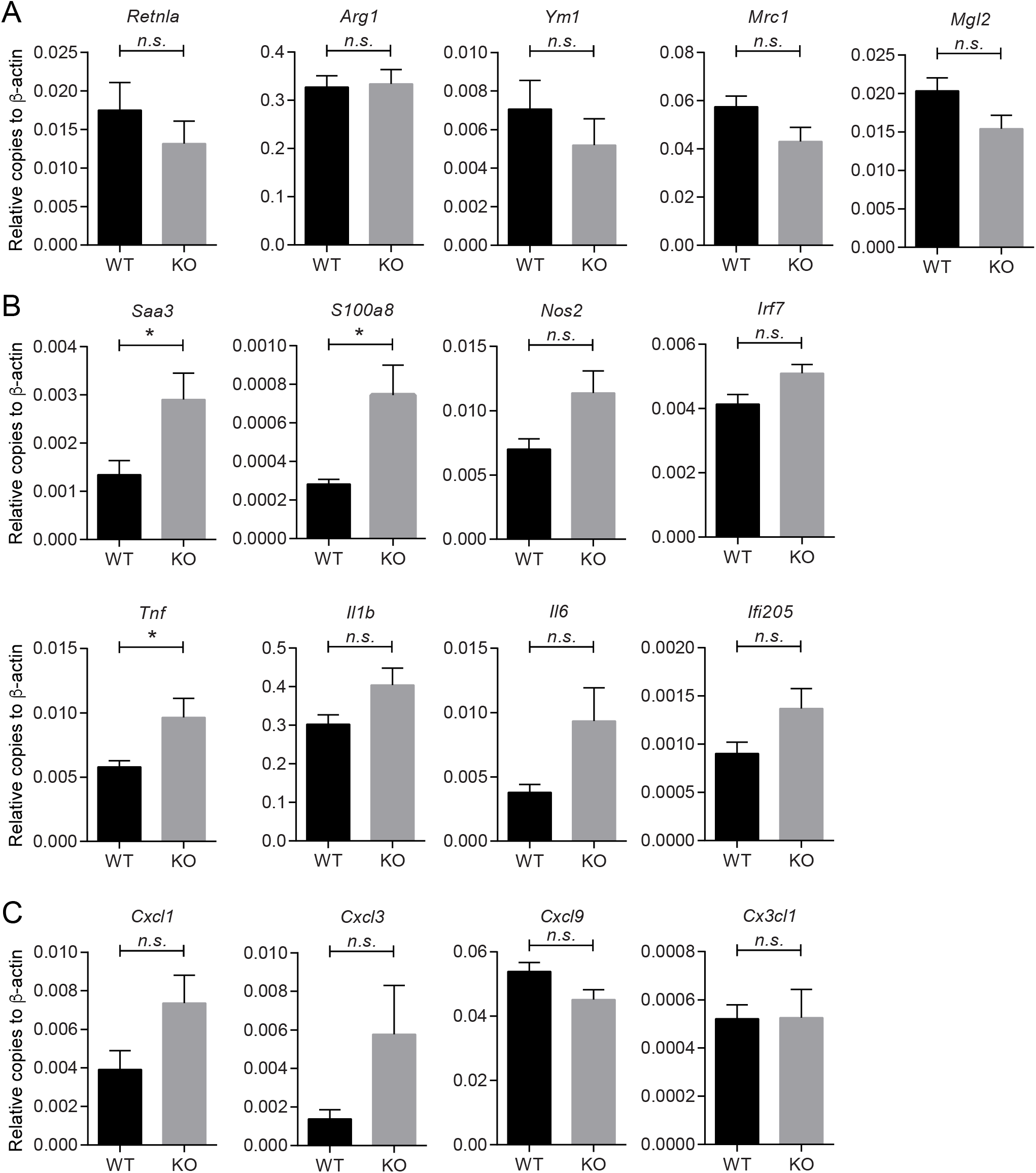
Impact of autophagy deficiency in myeloid cells on gene expression in TAMs from implanted tumors. Gene expression in TAMs from MC38 tumor at 21 days post implantation was further examined by quantitative RT-qPCR. WT indicates *Atg5*^flox/flox^ and KO indicates *Atg5*^flox/flox^*+LysMcre* mice. (A) Genes for alternatively activated macrophages. (B) Pro-inflammatory genes for classically activated macrophages. (C) Chemokines. n=4 per group. Error bars indicate mean +/- SEM, and statistical significance was calculated with unpaired T- test. *p < 0.05, n.s.: not significant.

**Figure S3.**
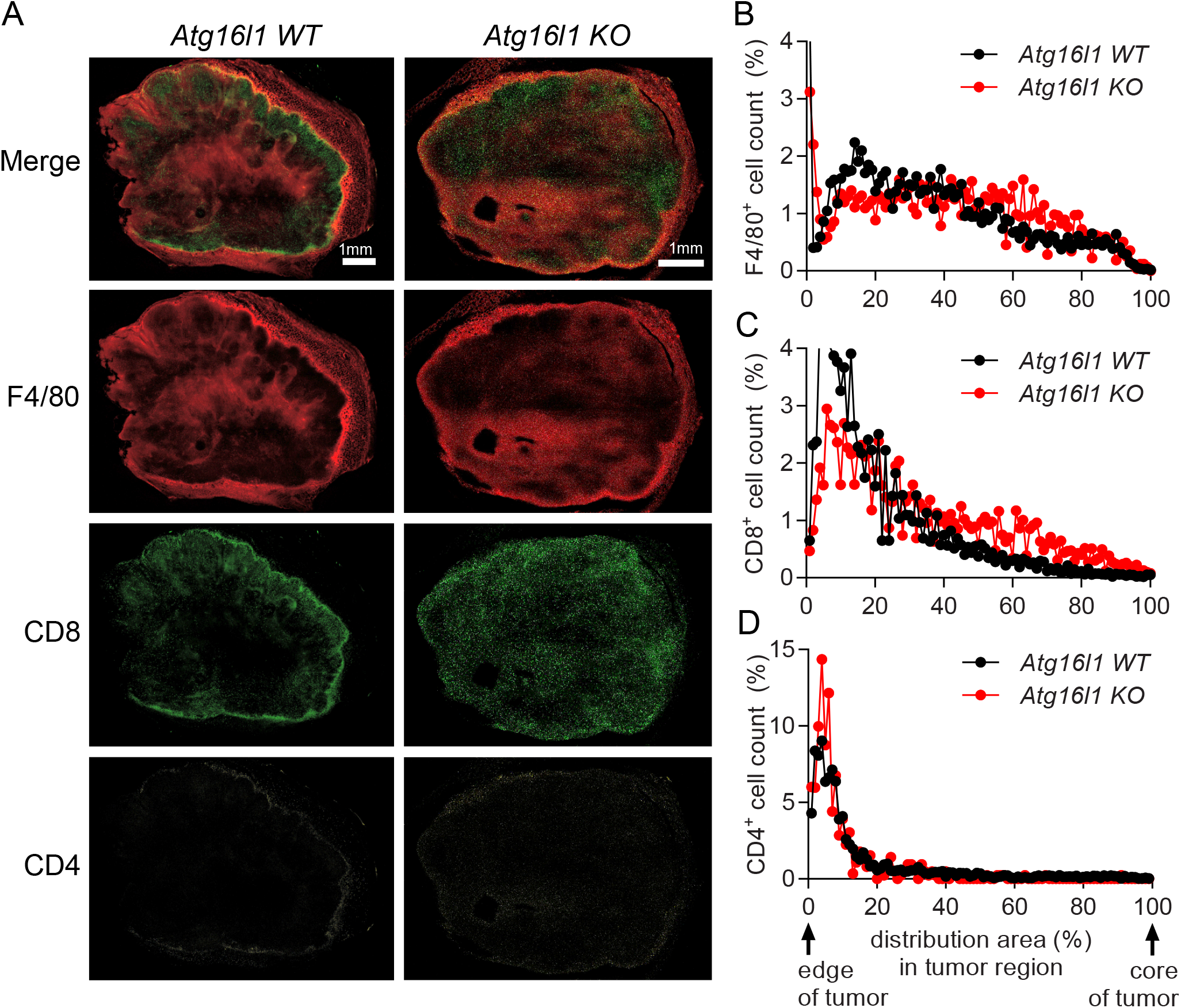
Autophagy deficiency in myeloid cells suppresses the growth of implanted tumors by affecting T cell infiltration. (A-D) Tumor tissues were collected at 21 days after 1x10^5^ MC38 cells were subcutaneously implanted in *Atg16l1 WT* (*Atg16l1*^flox/flox^) mice and *Atg16l1* KO (*Atg16l1*^flox/flox^*+LysMcre*) mice. n=3 per group. (A) Representative confocal microscopy images of the macrosections of tumors that were stained with anti-F4/80, anti-CD8 and anti-CD4 antibodies to visualize macrophages, CD8^+^ T cells and CD4^+^ T cells respectively. The distribution of immune cells, (B) F4/80^+^ cells, (C) CD8^+^ cells, and (D) CD4^+^ cells, in tumor sections was measured from the edge of the tumor to the core of the tumor.

